# HCFC1R1 Deficiency Blocks Herpes Simplex Virus-1 Infection by Inhibiting Nuclear Translocation of HCFC1 and VP16

**DOI:** 10.1101/2020.03.13.991679

**Authors:** Yangkun Shen, Zhoujie Ye, Xiangqian Zhao, Zhihua Feng, Jinfeng Chen, Lei Yang, Qi Chen

## Abstract

Upon HSV-1 infection, viral protein 16 (VP16), supported by Host Cell Factor C1 (HCFC1), is rapidly transported into the nucleus, and help to express a series of HSV-1 immediate-early proteins to begin its lytic replication. However, no direct evidence has shown if the HCFC1 deficiency can affect the proliferation of HSV-1 so far. Here, we showed that the HCFC1 deficiency led to a strong resistance to HSV-1 infection. Moreover, we identified Host Cell Factor C1 Regulator 1 (HCFC1R1) as a new host factor acting early in HSV infection for the transport of the HSV-1 capsid to the nucleus. The HCFC1R1 deficiency also led to a strong resistance to HSV-1 infection. The HCFC1R1 deficiency did not affect the attachment of HSV-1 to host cells but act early in HSV-1 infection by perturbing the formation of HCFC1/VP16 complex. Remarkably, in addition to wild-type HSV-1 infection, the host cells in the absence of either HCFC1 or HCFC1R1 showed strong resistant to the infection of TK-deficient HSV-1, which strain can course severe symptoms and tolerate to the current anti-HSV drug Acyclovir. Our data suggest that HCFC1 or HCFC1R1 may be used as the novel target for developing anti-HSV-1 therapies.

**IMPORTANCE:** Herpes simplex virus-1 (HSV-1) is widely spread in the human population and can cause a variety of herpetic diseases. Acyclovir, a guanosine analogue that targets the TK protein of HSV-1, is the first specific and selective anti-HSV-1 drug. However, the rapid emergence of resistant HSV-1 strains is occurring worldwide, endangering the efficacy of Acyclovir. Alternatively, targeting host factors is another strategy to stop HSV-1 infection. Unfortunately, although the HSV-1’s receptor, Nectin-1, was discovered in 1998, no effective antiviral drug to date has been developed by targeting Nectin-1. Targeting multiple pathways is the ultimate choice to prevent HSV-1 infection. Here we demonstrated that the deletion of HCFC1 or HCFC1R1 exhibits a strong inhibitory effect on both wild-type and TK-deficient HSV-1. Overall, we present evidence that HCFC1 or HCFC1R1 may be used as the novel target for developing anti-HSV-1 therapies with a defined mechanism of action.

---

Herpes simplex virus 1 (HSV-1) is a member of the alphaherpesvirus family, which can establish both lytic and latent infection (1, 2). Upon HSV-1 infection, HSV glycoprotein B or C mediates viral adhesion by binding to heparan sulfate proteoglycans on the cell surface, followed by HSV glycoprotein D interacting with Nectin-1, HVEM, and 3-O-sulfated heparan sulfate (3–5). For the viral genome that replicates in the nucleus, the viral entry also entails the extensive movement of viruses passing through the cytoplasm. In this context, the capsid of HSV-1 may be transported to the nuclear pore complex (NPC) through a vascular bundle since electronic microscopy analysis has shown neuronal microtubules bound by HSV-1 viral capsids (6–8).

HSV-1 expresses three groups of genes during its infection, including immediate-early (IE), early (E), and late (L) genes(9). These genes participate in various host’s regulatory pathways in assisting viruses to complete the HSV entire life cycle (10–12). HSV-1 proliferating process begins with the expression of immediate-early genes that contain the “TAATGARAT” motif, which is regulated by HSV-1 VP16 (13, 14). VP16 is a necessary factor for HSV-1 replication that needs to enter the nucleus(15). VP16 is dissociated from the viral capsid after HSV-1 infection, but it cannot enter the host cell nucleus autonomously and requires assistance from host cell factor C1 (HCFC1) which plays an important role as a member of the host cytokine family. HCFC1 is a diverse chromatin regulatory protein that is widely present in a variety of cells and regulates the cell cycle at various stages (16, 17). The C-terminus of HCFC1 contains two nuclear localization signal subunits, and the six β-helices at the N-terminus of HCFC1 are tightly bound together with the VP16 protein, which can bring VP16 into the nucleus upon their interaction (18, 19).

Upon moving into the nucleus, VP16 activates the expression of immediate early viral genes. However, VP16 does not directly recognize the “TAATGARAT” motif. VP16 is stabilized by forming a triplet complex with HCFC1 and Oct1 (20, 21). Upon binding, it immediately initiates transcription and translation of the viral immediate early genes, thereby creating the appropriate cell environment for replication and proliferation of HSV-1. Thus, HCFC1 is very important for early events in HSV infection, but no direct evidence so far as demonstrate that the HCFC1 deficiency can affect the proliferation of HSV-1. In addition, host cell factor C1 regulator 1 (HCFC1R1) can interact with HCFC1 and acts as its regulator(22). Overexpression of HCFC1R1 can cause a large accumulation of HCFC1 in the cytoplasm. However, it remains unclear whether HCFC1R1 can play a role in regulating HSV-1 infection.

Here, we demonstrate that either HCFC1 or HCFC1R1 deletion can inhibit HSV-1 proliferation. In addition, BGC823 cells in the absence of either HCFC1 or HCFC1R1 also showed strong resistance to the infection of thymidine kinase (TK) gene-deficient HSV-1 which strain can course severe symptoms and tolerate to the current anti-HSV drug Acyclovir. Furthermore, we show that the HCFC1/HCFC1R1 complex acts as an important scaffold to promote VP16 translocation into the nucleus. Our data suggest that HCFC1 and HCFC1R1 may be used as the new drug targets for anti-HSV-1 infection.

## RESULTS

### HCFC1 Is a Cellular Factor Required for HSV-1 Propagation

VP16 protein dissociates from the capsid and forms a complex with host cell factor 1 (HCFC1) after HSV-1 infects host cells. However, it remains no direct evidence if the HCFC1 deletion can lead to a resistant effect on HSV-1 proliferation. Therefore, we generated the HCFC1 knock out (KO) cell and examined the propagation of HSV-1 in the HCFC1 deficiency cell. We chose a gastric adenocarcinoma cell line BGC-823, which is sensitive to the infection of HSV-1. Firstly, we established a clonal cell line null for HCFC1 in BGC823 cells using the CRISPR/Cas9 system (Figure 1A). We identified three HCFC1-KO cell monoclones and Western blotting results showed absence of the HCFC1 protein (Figure 1B). We analyzed the genome and amino acid sequences of the third HCFC1-KO cell strain and sequencing data revealed a frameshift mutation in the knockout cells, indicating the successful construction of BGC823^HCFC1−/−^ cell line (Figure 1C). To visually observe the virus proliferation in the host cell, we generated a fluorescent HSV-1 virus, referred to as HSV-1-VP26-mCherry virus, in which a red fluorescent protein gene is in frame fused after the VP26 gene (Figure 1D). When infected with the HSV-1-VP26-mCherry virus (MOI=1), BGC823 cells were luminous, indicating successful construction of the fluorescent HSV-1-VP26-mCherry virus (Figure 1E). Furthermore, the cell death rate induced by HSV-1-VP26-mCherry was equivalent to wild type HSV-1 virus (Figure 1F), suggesting that the recombinant fluorescent fusion protein with VP26 did not affect the assembly and infectivity of the HSV-1 virus.

**Figure 1.**
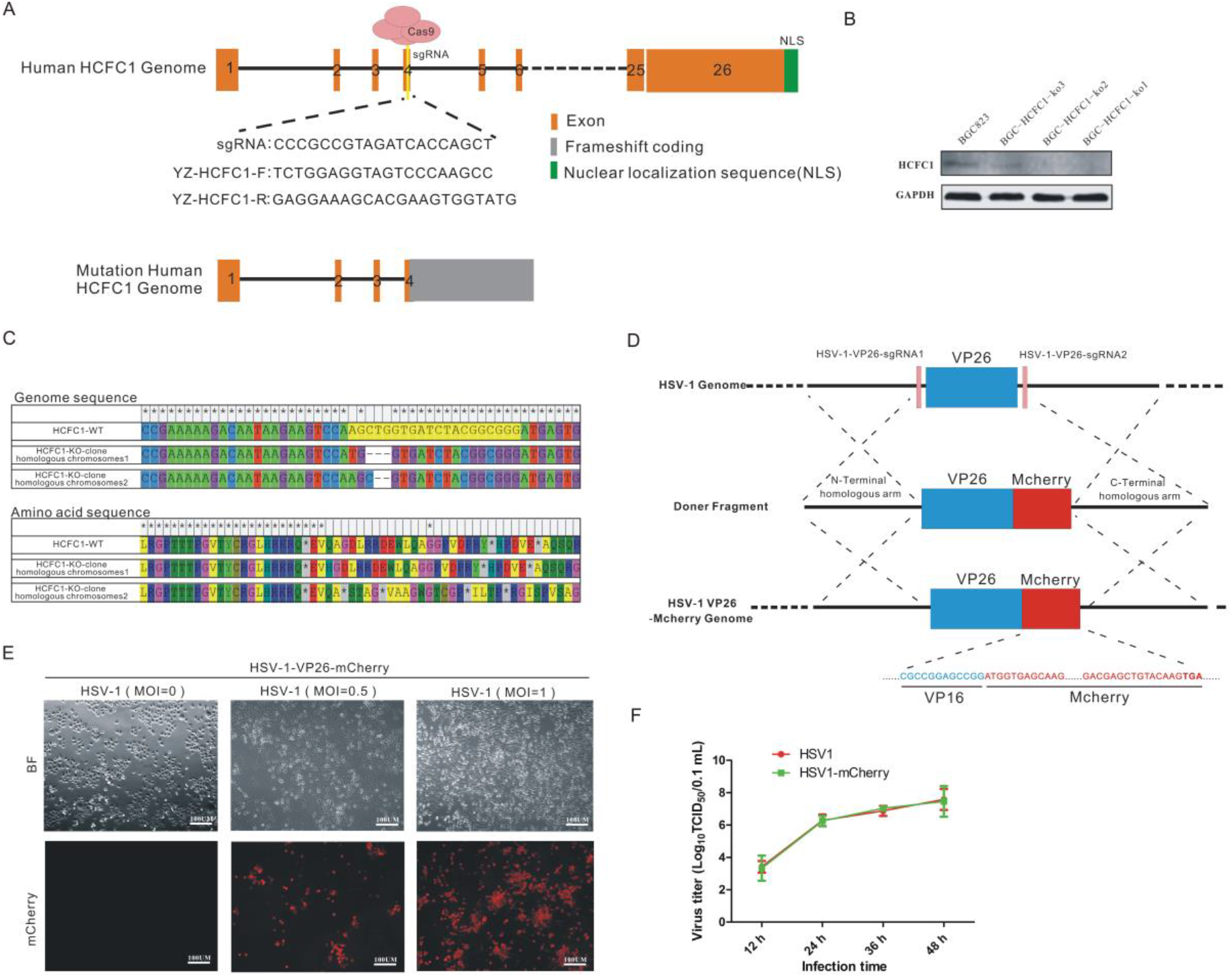
Construction of the BGC-823^HCFC1−/−^ knockout cell line and HSV-1-VP26-mCherry virus. (A) The schematic diagram of HCFC1 wild-type (WT) and knockout (KO) gene alleles. (B) Identification of the HCFC1 knockout cell line by Western blotting. WT and HCFC1 KO BGC-823 cells were detected by anti-HCFC1 antibody. (C) Sequencing analysis of the BGC-823 KO cell line at the HCFC1 genomic locus. The genome DNA was sequenced using a pair of primers flanking the HCFC1 gene. The cell genotype was identified by T-A cloning and sequencing; The gene deletion and changes of corresponding amino acid coding are indicated. (D) Schematic diagram showing the construction of the fluorescent HSV-1 virus, which fuses a red fluorescent protein gene behind the VP26 gene. (E) Infection of HSV-1-VP26-mcherry virus (red) causes BGC-823 cell death. Cells were plated with a lower cell density (10^5^/6 well plate), and infected with HSV-1 (MOI = 0.5, 1) after 48 hour. Images were collected by Zeiss microscope (Oberkochen, Germany). Changes in cell death during time points are shown. Scale bar, 100 μm. (F) WT and HSV-1-VP26-mcherry virus had the same infection ability. BGC-823 cells were infected with HSV-1 and HSV-1-VP26-mcherry virus (MOI=1), respectively for 2 hour, washed twice with PBS, and replaced with fresh medium. Cells together with the supernatants were harvested at 12, 24, 36 and 48 hour after infection. After three repeated freeze-thaw cycles, the samples were centrifuged at 10,000 rpm for 5 min, and then the viral titer determined by a standard viral plaque assay using Vero cells. pfu, plaque-forming units.

We then used the HSV-1-VP26-mCherry virus to monitor the infection of HSV-1 in the wild-type (WT) and HCFC1 deficiency BGC-823 cells. As expected, BGC823^HCFC1−/−^ cells showed strong resistance to the virus (Figure 2A). And the infection of HSV-1 for 24 to 72 hours led to the cell loss in WT but not HCFC1-deficient BGC-823 cells (Figure 2B). Furthermore, the titer of HSV-1 produced in HCFC1 KO cells was markedly lower than that in WT BGC-823 cells (Figure 2C). We further examined the cell viability following the viral infection, and the data again showed that the deletion of HCFC1 could effectively cause resistance to HSV-1 infection (Figure 2D). These data suggest that HCFC1 is a host factor that requires for HSV-1 propagation.

**Figure 2.**
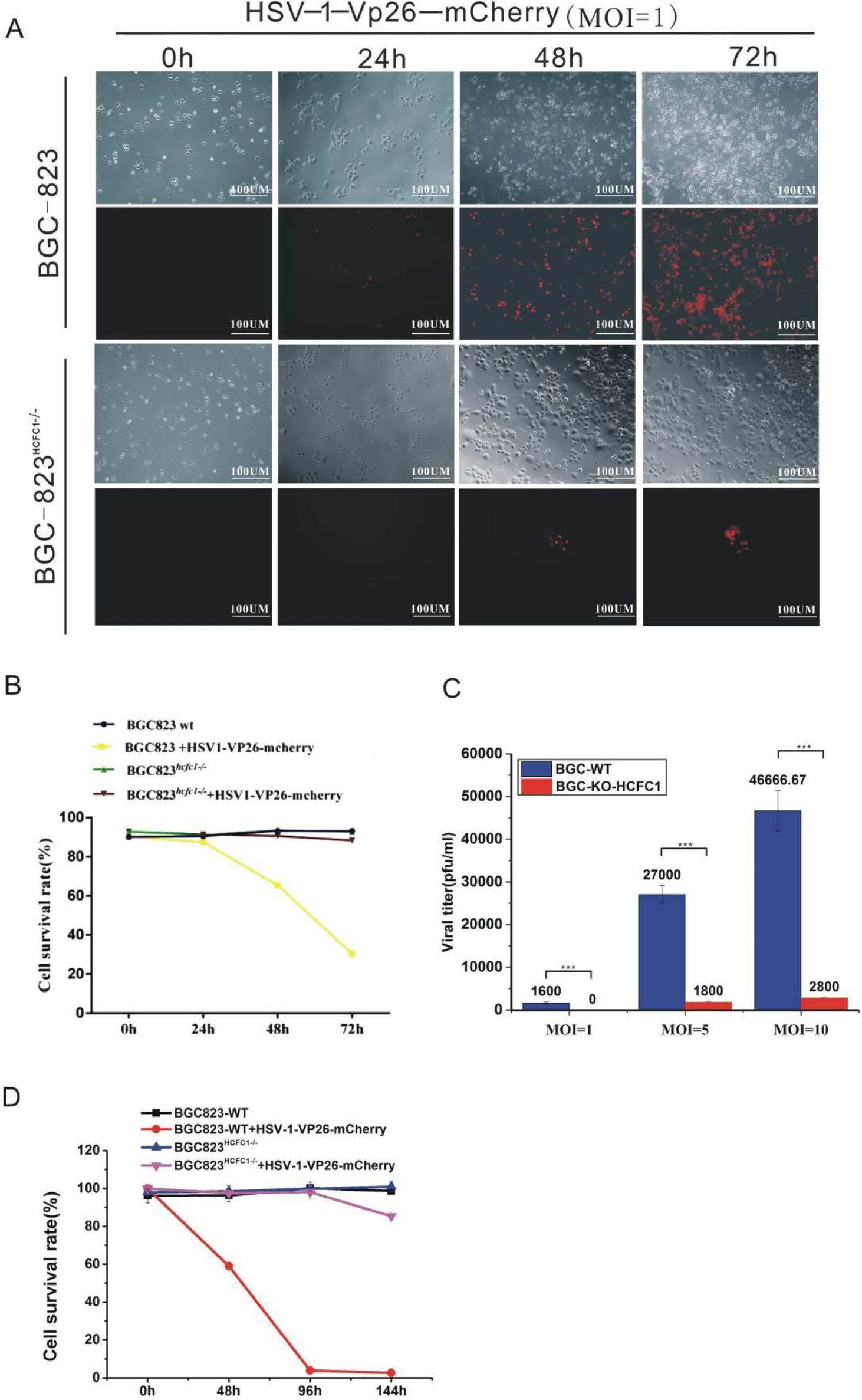
Antiviral analyses of the HCFC1 gene. (A) HCFC1 KO BGC-823 cells are resistant to HSV-1-VP26-mcherry infection compared with WT BGC-823 cells. Time-lapse (0, 24, 48, 72 hours) images of WT and HCFC1 KO BGC-823 upon HSV-1-VP26-mcherry (MOI = 1) (red) infection using Zeiss microscope. Changes in cell death during time points are shown. Scale bar, 100μm. (B) Changes of cell survival rate following HSV-1 infection in WT and HCFC1 KO BGC-823 cells. Cells were seeded in 6 cm cell culture plates at a 10^5^ cell density. After 24 hours, cells were infected with HSV-1 (MOI = 1). Cell viability was determined by trypan blue staining at different time points (0, 24, 48, 72 hours). Data are represented as mean ± SD. (C) HCFC1 deficiency restricted HSV-1 propagation. WT and HCFC1 KO BGC-823 cells were infected with HSV-1 (MOI=0, 1, 5, 10) for 2 hr, washed twice with PBS, and replaced with fresh medium. Cells together with the supernatants were harvested at 24 hour after infection. After three repeated freeze-thaw cycles, the samples were centrifuged at 10,000 rpm for 5 min, and then the viral titer was determined by a standard viral plaque assay using Vero cells. pfu, plaque-forming units. (D) The resistance of WT and HCFC1 KO BGC-823 cells to HSV-1 viruses following the time course. Cells were seeded in 6 cm cell culture plates at a 10^5^ cell density. After 24 hours, cells were infected with HSV-1 (MOI = 1). Cell viability was determined by trypan blue staining at different time points (0, 48, 96, 144 hours). Data are presented as mean ± SD.

### HSV-1 Infection Leads to HCFC1-Dependent Nuclear Localization of VP16

Since VP16 needs to be localized to the nucleus to initiate the replication of the HSV-1 genome, we first investigated the cellular localization of VP16. We found that VP16 was localized in the nucleus when overexpressed alone (Figure 3A). Interestingly, fluorescence microscopy data showed that VP16 was located not only inside the nucleus, but also in the surrounding of the nucleus (Figure 3B). We supposed that, because of the colocalization of HSV-1 and VP16, most of the VP16 proteins might be gathered in the nuclear pore complex (NPC) for HSV-1 assembly. Since VP16 can form a complex with HCFC1 (23, 24), we asked if HCFC1 was needed for VP16 translocation to the nucleus. We overexpressed VP16 in BGC823^HCFC1−/−^ cells, and found that the deletion of HCFC1 blocked the nuclear localization of VP16 (Figure 3C). These data suggest that HCFC1 is required for promoting VP16 entry into the nucleus.

**Figure 3.**
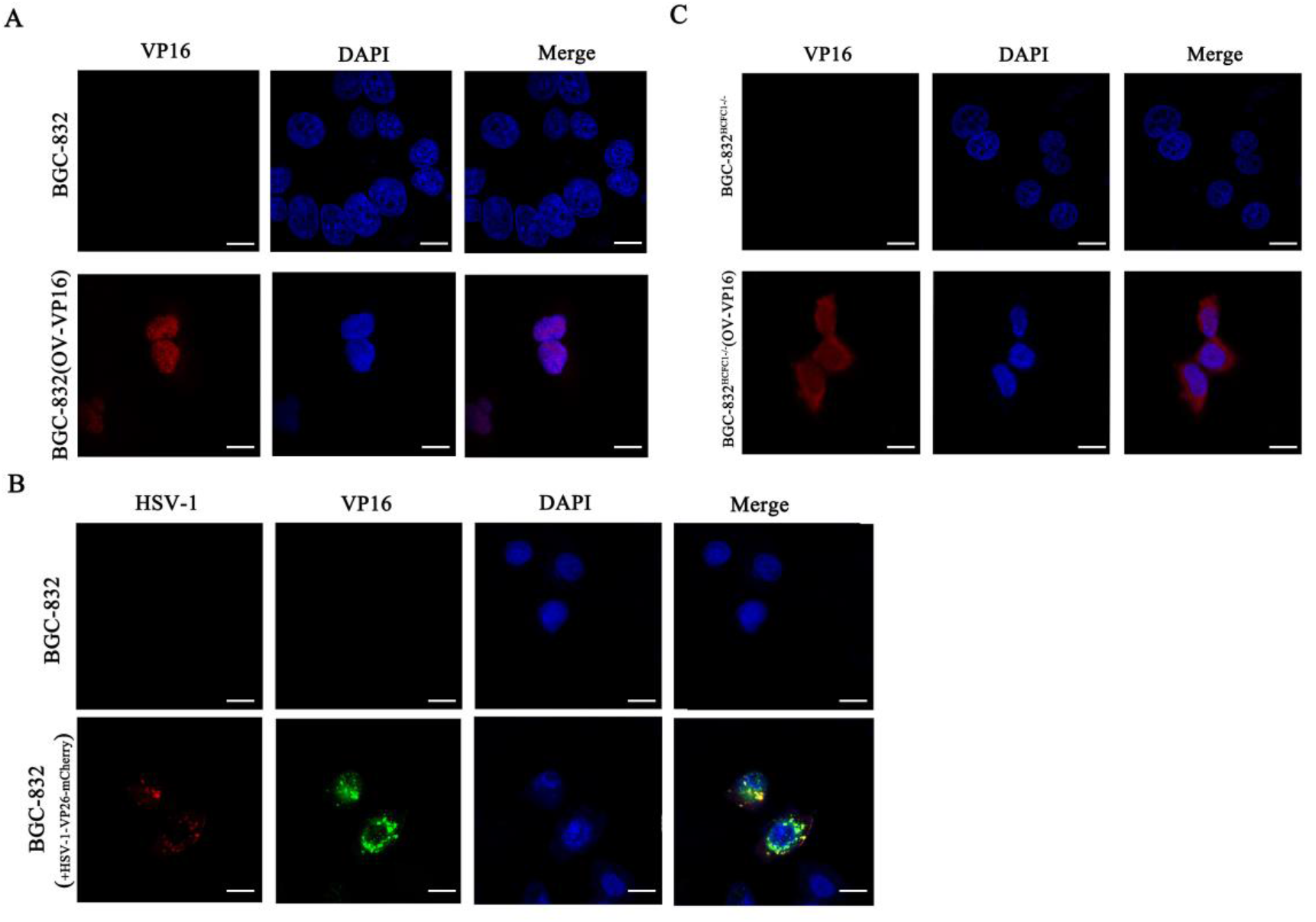
HSV-1 infection leads to HCFC1-dependent nuclear localization of VP16. (A) VP16 could be guided into the nucleus in support of host factors. Cells were seeded in confocal dish at a 5×10^4^ cell density. After 12 hours, cell transfection was done with OV-FLAG-VP16 vector (2 μg) using Lipofectamine 3000. Verification of subcellular localization of VP16 in BGC-823 cells 24 hour after transfection using anti-VP16 antibody (red) and DAPI (Nuclear marker, blue). Scale bar, 20μm. (B) Fluorescence microscopy showed that VP16 was localized in the nucleus after HSV-1 infection. Cells were seeded in confocal dish at a 5×10^4^ cell density. After 12 hours, cells were infected with HSV-1-VP26-mcherry (MOI=1) (red) for 2 hour, washed twice with PBS, and replaced with fresh medium. Verification of subcellular localization of VP16 in BGC-823 cells 24 hour after transfection using anti-VP16 antibody (green) and DAPI (blue). Scale bar, 20μm. (C) HCFC1 deficiency prevents VP16 (red) getting into the nucleus. The same as in (A), except HCFC1 KO cells were used. Scale bar, 20μm.

### HCFC1R1 Is Required for the Transport of HSV Capsid to the Nucleus

HCFC1R1 is an interaction protein of HCFC1, and regulates HCFC1 activity by modulating its subcellular localization. Therefore, we asked if HCFC1R1 played any role in promoting VP16 entry into the nucleus. We generated a HCFC1R1 knockout BGC823 cell line (BGC823^HCFC1R1−/−^) using the CRISPR/Cas9 system (Figure 4A). We identified three HCFC1R1 KO cell monoclones and Western blotting data showed the absence of HCFC1R1 protein in these cell clones (Figure 1B). The sequencing data revealed a frameshift mutation in the HCFC1R1 KO cells (Figure 4C).

**Figure 4.**
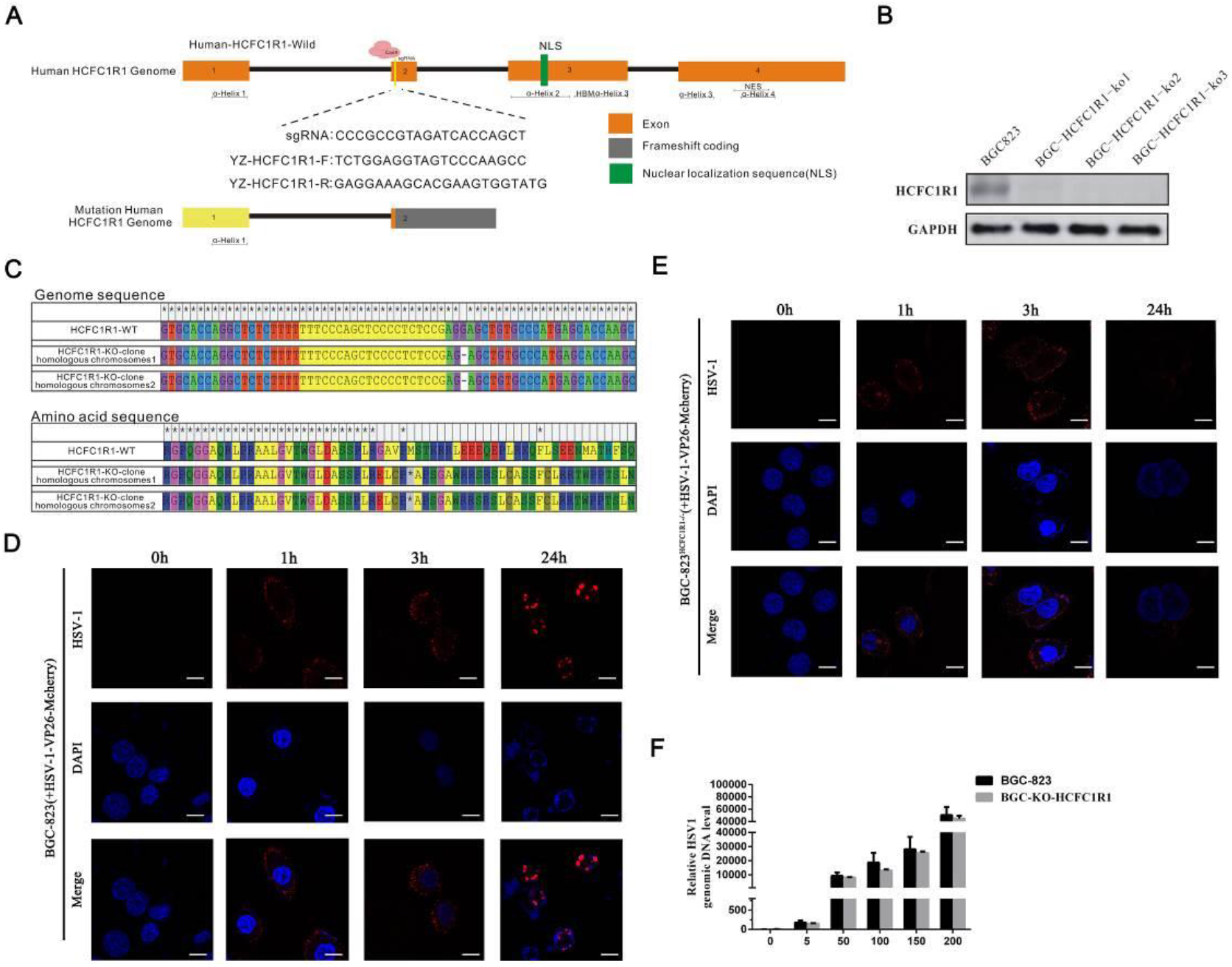
HCFC1R1 is a host factor required for the transport of HSV capsid to the nucleus. (A) Schematic diagram of HCFC1R1 WT and KO gene alleles. (B) Identification of the HCFC1R1 knockout cell line by Western blotting using anti-HCFC1R1 antibody. (C) Sequencing data of the BGC-823 knockout cell line at the HCFC1R1 genomic locus. The genome DNA was sequenced using a pair of primers flanking the HCFC1R1 gene. Genotyping was done by T-A cloning and sequencing, the gene mutation in HCFC1R1 KO cells was indicated. (D) Immunofluorescence showing the viral infection process in WT BGC-823 cells. Cells were plated at a 10^4^ cell density, and infected with HSV-1-VP26-mcherry (MOI = 100) for 12 hour. Photos were collected at the different infection time points (1, 3, 24 hours). The HSV-1-VP26-mcherry viruses accumulated in the cell membrane at 1 hour after infection, gradually in the cytoplasm at 3 hours, and eventually forming the punctate structure in the nucleus at 24 hours after infection. Scale bar, 20μm. (E) The same as in (D), except HCFC1R1 KO cells were used. However, the process of aggregation of HCFC1 into the nucleus was not seen after HSV-1 infection in HCFC1R1 deficiency BGC823 cells. Scale bar, 20μm. (F) Detection of the ability of viral binding using a quantitative binding assay. WT and HCFC1R1 KO BGC-823 cells were seeded in 6 cm cell culture plates at a 10^5^ cell density. After 12 hours, cells were treated with various titers of HSV-1 (MOI = 0, 5, 50, 100, 200) for 1 hour at 37°C. The viral genome was extracted from the cells. The copy number of HSV-1 binding was examined by qPCR. (***, *p* < *0.001*; **, *p* < *0.01*;*, *p* < *0.05*). Data are presented as mean ± SD.

To understand if HCFC1R1 affects HSV-1 infection, we infected wild-type and HCFC1R1 deficient BGC823 cells with the HSV-1-VP26-mCherry virus. Immunofluorescence data showed that when infected wild-type BGC823 cells with a high titer virus, a large number of HSV-1 accumulated around the cell membrane at 1 hour, gradually in the cytoplasm at 3 hours, and eventually forming a punctate structure in the nucleus at 24 hours (Figure 4D). In BGC823^HCFC1R1−/−^ cells, a large number of HSV-1 were also accumulated around the cell membrane at 1 hour. The deletion of HCFC1R1 affected the spread of the HSV-1 virus at 3 hours. Strikingly, the accumulation of HCFC1 in the nucleus was not seen in BGC823^HCFC1R1−/−^ cells (Figure 4E), and the virus particles had disappeared at 24 hours, suggesting the HCFC1R1 deficiency leads to failure of the propagation of HSV-1.

To ask if HCFC1R1 was required for HSV-1 binding, we assessed the content of HSV-1 on the cell membrane using a quantitative binding assay and found that loss of HCFC1R1 did not block HSV-1 binding to the host cell (Figure 4F). These data suggest that the HCFC1R1 deficiency does not affect the virus binding to the plasma membrane, but rather prevents the virus from entering the nucleus.

### The HCFC1R1 Deficiency Inhibits HSV-1 Propagation

We next examined the effect of HSV-1 infection in BGC823^HCFC1R1−/−^ cells. Our data showed that the cell viability of WT BGC823 cells gradually decreased following the propagation of viral infection, while the cell viability of BGC823^HCFC1R1−/−^ was much less affected by HSV-1 infection (Figure 5A). Furthermore, the titer of HSV-1 produced in BGC823^HCFC1R1−/−^ cells was markedly lower than that in WT BGC-823 cells after HSV-1 infection (Figure 5B). When infected with HSV-1-VP26-mCherry virus (MOI =1), the number of red fluorescent BGC823^HCFC1R1−/−^ cells was significantly lower than the number of WT BGC823 cells after 48 h infection (Figure 5C). BGC823^HCFC1R1−/−^ cells kept in a good growth state regardless of HSV-1-VP26-mCherry virus infection while WT BGC823 cells gradually died following the infection time course.

**Figure 5.**
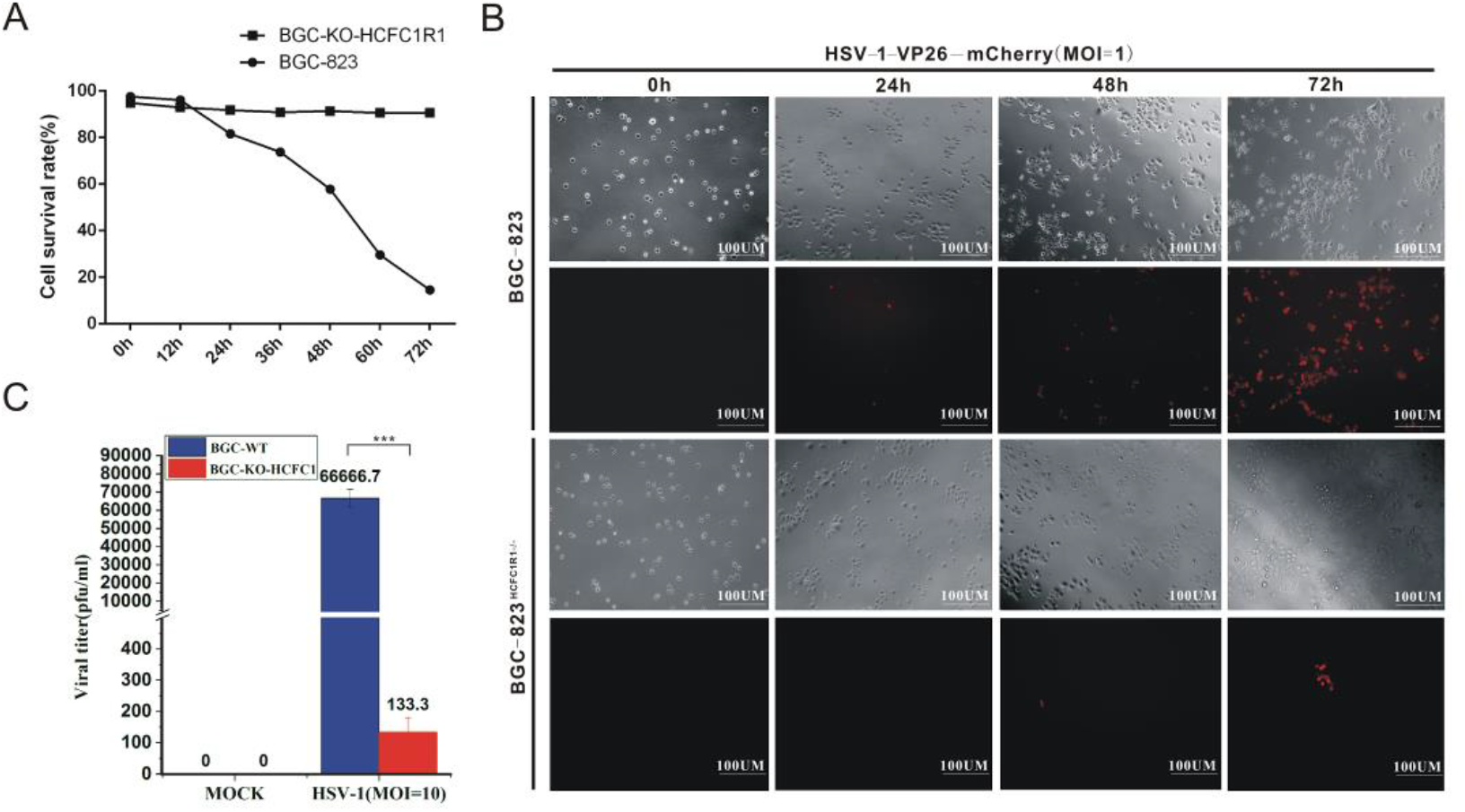
HCFC1R1 knockout inhibits HSV infection. (A) Changes of cell survival rate following HSV-1 infection in WT and HCFC1R1 KO BGC-823 cells. Cells were seeded in 6 cm cell culture plates at a 10^5^ cell density. After 24 hours, cells were infected with HSV-1 (MOI = 1). Cell viability was determined by trypan blue staining at different time points (0, 12, 24, 36, 48, 60, 72 hours). Data are presented as mean ± SD. (B) HCFC1R1 KO BGC-823 cells are resistant to HSV-1-VP26-mcherry infection compared with WT BGC-823 cells. The cells were infected with HSV-1-VP26-mcherry (MOI=1) for 24, 48, 72 hours, and then the antiviral effect was assessed by using Zeiss fluorescence microscope. Changes in cell death during time points are shown. Scale bar, 100 μm. (C) HCFC1R1 deficiency restricted HSV-1 propagation. WT and HCFC1 KO BGC-823 cells were infected with HSV-1 (MOI=1) for 2 hour, washed twice with PBS, and replaced with fresh medium. Cells together with the supernatants were harvested at 24 hour after infection. After three repeated freeze-thaw cycles, the samples were centrifuged at 10,000 rpm for 5 min, and then the viral titer was determined by a standard viral plaque assay using Vero cells. pfu, plaque-forming units.

HSV-1 expresses immediate early, early, and late genes at the various infection stages to complete its entire life cycle. The HSV- gB gene is expressed through the entire HSV-1 life cycle. To test if HCFC1R1 affected the viral proliferation, we examined the HSV-1 genome copies by examining the HSV-1 gB gene expression in BGC-823^HCFC1R1−/−^ cells. The HSV-1 gB gene copy number was clearly reduced as shown by qPCR, suggesting that the HCFC1R1 deficiency affects HSV-1 propagation (Figure 6A). To further ask if loss of HCFC1R1 was correlated with the life cycle of HSV-1, we analyzed the mRNA expression of several viral genes including ICP0 (representing immediate early gene), TK (early gene), and GD (late gene) in the control and BGC-823^HCFC1R1−/−^ cells by qRT-PCR in 24, 48, and 72 hours after infection (25–27). qRT-PCR data showed that missing HCFC1R1 significantly inhibited the mRNA transcription in BGC-823^HCFC1R1−/−^ cells at the various time points (Figure 6B, C, D). Western blotting data also showed that the levels of viral proteins were reduced in BGC-823^HCFC1R1−/−^ cells relative to the control (Figure 6E). Therefore, we suppose that the significant low expression of HSV-1 viral genes correlates with the decrease in the replication and proliferation of HSV-1 in BGC823^HCFC1R1−/−^ cells, and suggest that the HCFC1R1 gene deletion results in the inability of the HSV-1 virus to replicate normally in the host cell.

**Figure 6.**
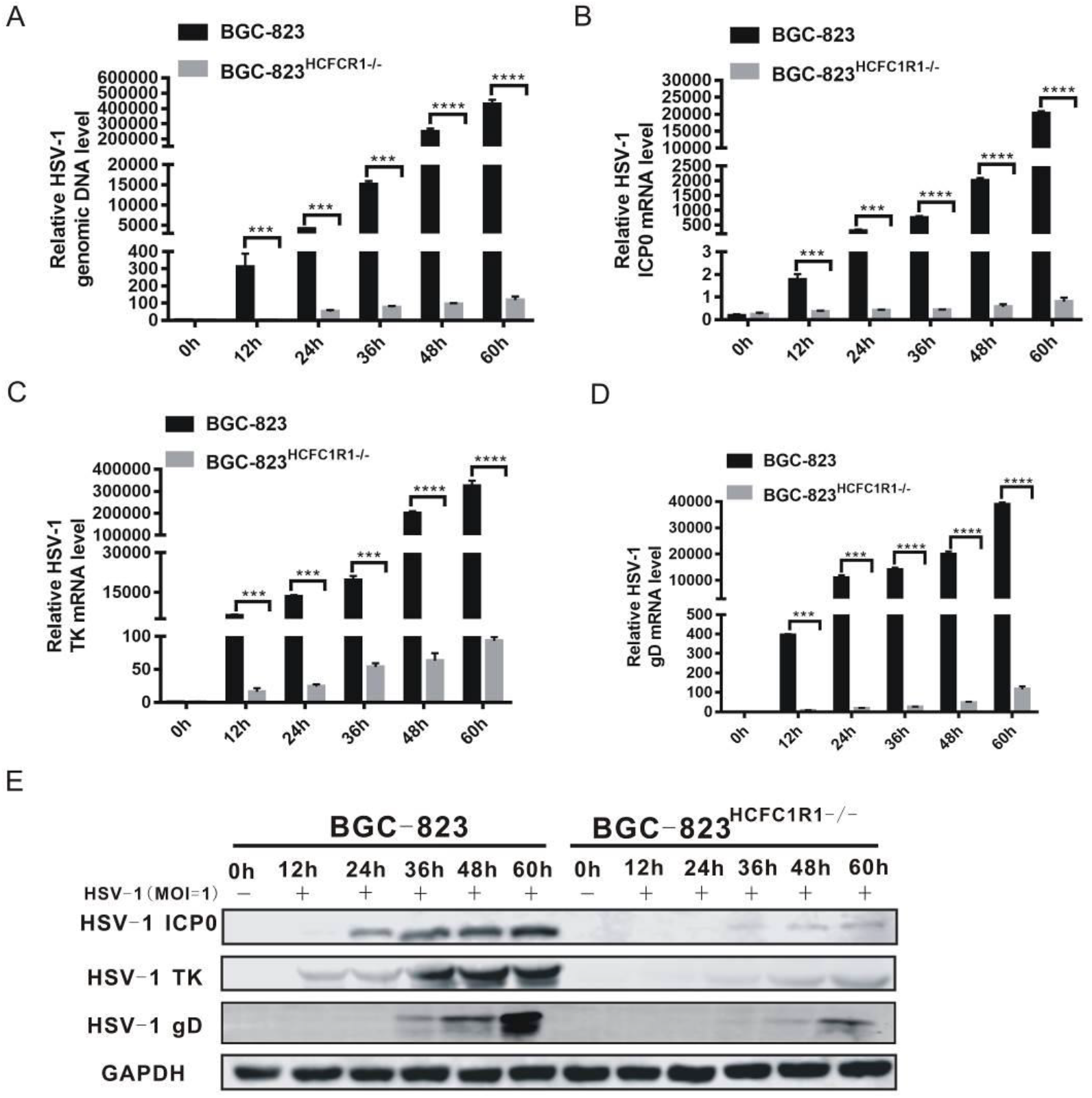
HCFC1R1 knockout restricts HSV-1 propagation. (A) Quantification of the relative genome copy numbers of HSV-1 by qPCR at different time points after viral infection (HSV-1 GD gene was used to represent). (***, *p* < *0.001*; **, *p* < *0.01*; *, *p* < *0.05*). Data are presented as mean ± SD. (B-D) Quantification of the relative mRNA expression of HSV-1 ICP0, TK and GD by qRT-PCR. The data showed that the BGC-823^HCFC1R1−/−^ cells were resistant to HSV-1 infection, while WT BGC-823 cells died gradually with the increase of time. (***, *p* < *0.001*; **, *p* < *0.01*;*, *p* < *0.05*). Data are presented as mean ± SD. (E) Detection of the expression of ICP0, TK and GD proteins after viral infection for different time points using Western blotting in BGC cells expressing HCFC1R1 or not.

### HCFC1R1 Is Required for the Formation of HCFC1 and VP16 Complex

Since both the viral VP16 protein and the host HCFC1R1 are required for HSV-1 to enter the nucleus, it is possible that HCFC1R1 may affect HSV-1 migration by regulating VP16 transportation. To test this possibility, we overexpressed VP16 and examined its localization in BGC823^HCFC1R1−/−^ cells. We found that in contrast to the control cells, VP16 was hardly localized in the nucleus of BGC823^HCFC1R1−/−^ cells (Figure 7A), suggesting that the HCFC1R1 deficiency can affect the transport of VP16 to the nucleus. Previous studies have shown that HCFC1 can affect the entry of VP16 proteins into the nucleus, thus affecting the immediate early gene expression of the virus. Therefore, we asked if HCFC1R1 could affect the migration of VP16 through regulating the entry of HCFC1 into the nucleus. We examined the HCFC1 localization in the host cell and found that the HCFC1 entry to the nucleus was significantly impaired in HCFC1R1 deficiency cells (Figure 7B). Thus, our data suggest that HCFC1R1 may affect VP16 proteins entering into the nucleus by regulating the HCFC1 activity.

**Figure 7.**
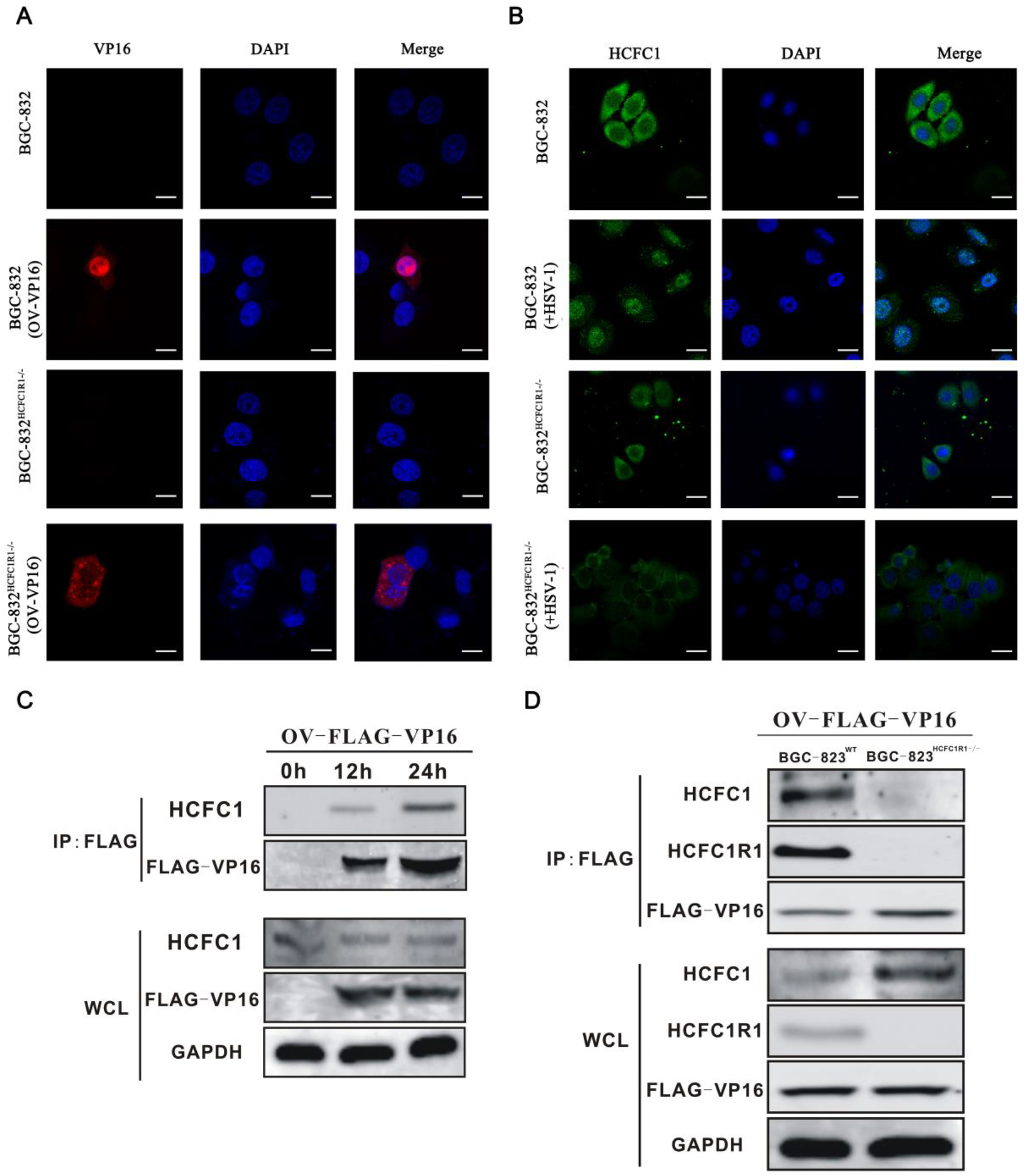
HCFC1R1 regulates the formation of HCFC1 and VP16 complex. (A) Subcellular localization of VP16 in WT and HCFC1R1-KO BGC-823 cells. HCFC1R1 deficiency prevents VP16 getting into the nucleus. Cells were seeded in confocal dish at a 5×10^4^ cell density for 12 hours and transfected with OV-FLAG-VP16 vector (2 μg) using Lipofectamine 3000. The cells were collected 24 hour after transfection, and stained by anti-VP16 antibody (red) and DAPI (blue). Scale bar, 20μm. (B) HCFC1R1 deficiency prevents HCFC1 getting into the nucleus. Cells were seeded in confocal dish at 5×10^4^ for 12 hours, and infected with wild-type HSV-1 (MOI=1) for 2 hour. Subcellular localization of HCFC1 in WT and HCFC1R1 KO BGC-823 cells was analyzed by using anti-HCFC1 antibody (green) and DAPI (blue). Scale bar, 20μm. (C) HSV-1 infection leads to the formation of HCFC1 and VP16 complexe. Cells were seeded in 15 cm plates at a 1×10^7^ cell density. After 12 hours, the cells were transfected with the OV-FLAG-VP16 vector (20ug) using Lipofectamine 3000 for 12 or 24 hour. The cell lysates were immunoprecipitated with anti-Flag beads, the interaction of HCFC1 and VP16 was determined by immunoblotting with anti-HCFC1 and anti-FLAG antibodies. (C) HCFC1R1 deficiency prevents the formation of HCFC1 and VP16 complex. The same as in (C), except HCFC1R1 deficiency cells and anti-HCFC1R1 antibody were used.

VP16 can form a complex with HCFC1, which brings VP16 entering the nucleus. Therefore, we asked if HCFC1R1 could affect the formation of the HCFC1 and VP16 complex. We overexpressed Flag-VP16 in the wild-type and HCFC1R1 deficiency BGC-823 cells and analyzed the protein interactions between HCFC1 and VP16. Co-immunoprecipitation data showed that the formation of the HCFC1 and VP16 complex was induced by HSV-1 infection, but was suppressed by the HCFC1R1 deficiency (Figure 7C-D). Furthermore, HCFC1R1 was also able to directly interact with HCFC1 and VP16 (Figure 7). Taken together, these data suggest that HCFC1R1 may affect VP16 entering the nucleus by regulating the HCFC1 complex formation.

### Comparison of Anti-HSV-1 Efficiency by Targeting HCFC1 or HCFC1R1 with by Targeting Nectin-1

The receptors are one of the most important targets for developing antiviral drugs. Nectin-1 is considered as a major receptor for HSV-1 (28, 29). Therefore, we assessed the anti-HSV-1 efficiency by targeting HCFC1 or HCFC1R1 in comparison with by targeting Nectin-1. We generated Nectin-1-knockout cells and infected these cells with HSV-1 and found that the HSV-1 infection caused the WT cell loss but had little effect on Nectin-1-KO BGC-823 cells (Figure 8A), suggesting that the Nectin-1 knockout cells are resistant to HSV-1 infection (Figure 8B). Next, we compared the antiviral effect of HCFC1-KO and Nectin-1-KO cells, and found that the anti-HSV1 efficiency was comparable in both these KO-cell lines (Figure 8C). Consistently, HCFC1R1 knockout cells also showed a similar effect in blocking HSV-1 infection (Figure 8D).

**Figure 8.**
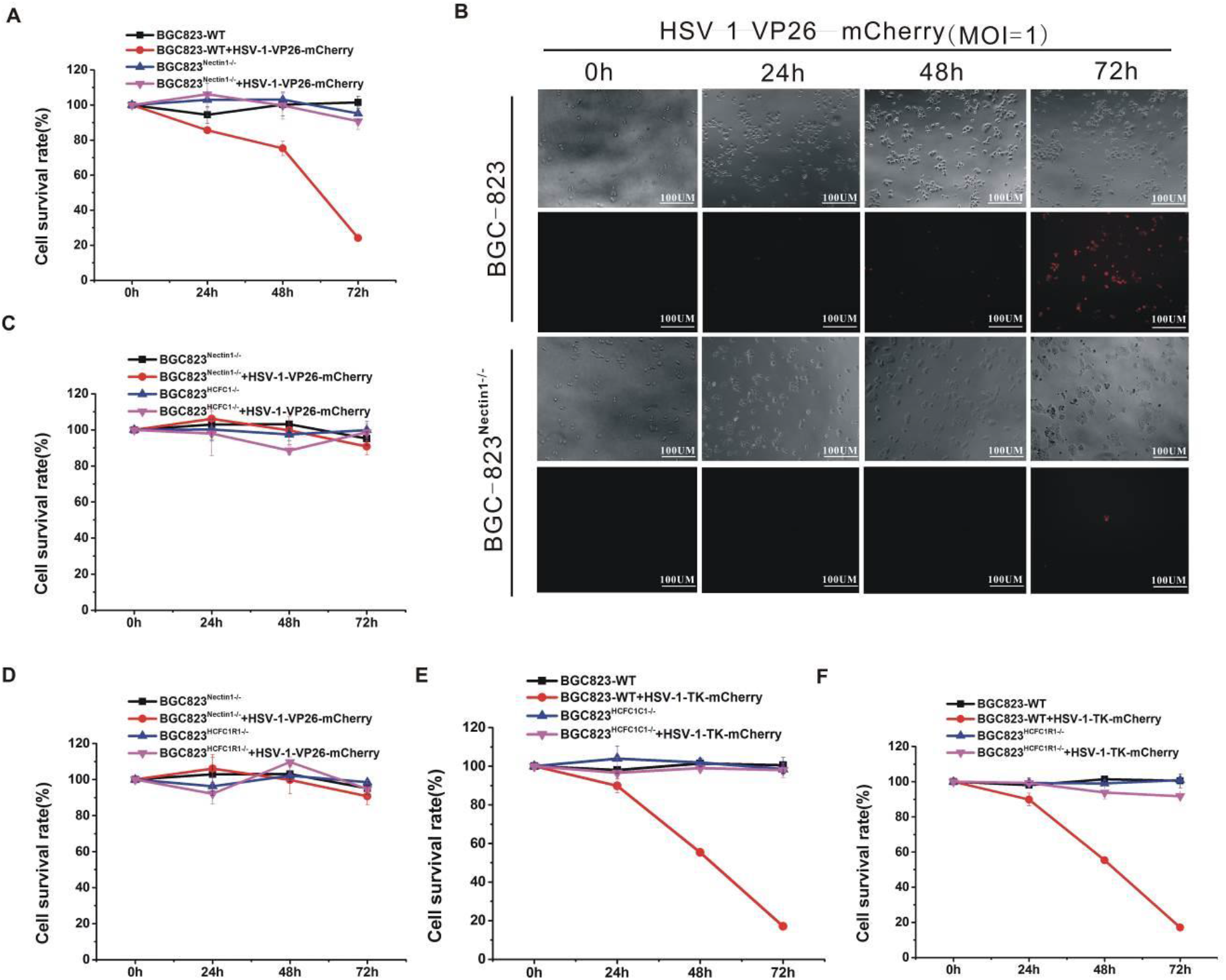
Deletion of HCFC1 or HCFC1R1 has a strong inhibitory effect on the TK-deficient HSV-1. (A) Changes of cell survival rate following HSV-1 infection in WT and HCFC1 KO BGC-823 cells. Cells were seeded in 6 cm cell culture plates at a 10^5^ cell density. After 24 hours, cells were infected with HSV-1 (MOI = 1). Cell viability was determined by trypan blue staining at different time points (0, 24, 48, 72 hours). Data are presented as mean ± SD. (B) HCFC1 KO BGC-823 cells are resistant to HSV-1-VP26-mcherry (red) infection compared with WT BGC-823 cells. The antiviral effect was assessed by cell death using Zeiss microscope. Cells were plated with a 10^5^ density, and infected with HSV-1 (MOI = 1) for 12 hour. Images were collected at the different time points (24, 48, 72 hours). Changes in cell death during time points are shown. Scale bar, 100 μm. (C) HCFC1 and Nectin1 knockout cells have the same resistance to HSV-1. Cells were seeded in 6 cm cell culture plates at a 10^5^ cell density. After 24 hours, cells were infected with HSV-1 (MOI = 1). Cell viability was determined by trypan blue staining at different time points (0, 24, 48, 72 hours). Data are presented as mean ± SD. (D) The same as in (C), except HCFC1R1 KO cells were used. (E) Changes of cell survival rate following HSV-1-KO-TK infection in WT and HCFC1 KO BGC-823 cells. Cells were seeded in 6 cm cell culture plates at a 10^5^ cell density. After 24 hours, cells were infected with HSV-1-KO-TK (MOI = 1). Cell viability was determined by trypan blue staining at different time points (0, 24, 48, 72hours). Data are presented as mean ± SD. (F) The same as in (D), except HCFC1R1 KO cells were used.

### Deletion of HCFC1 or HCFC1R1 Has A Strong Inhibitory Effect on the TK-Deficient HSV-1

Currently, Acyclovir and its analogues have been used for preventing and treating HSV-1 infection (30–32). However, the TK-deficiency disables the Acyclovir antiviral effects and currently no therapy for TK-deficient HSV-1 has been developed. Therefore, we asked if targeting either HCFC1 or HCFC1R1 could affect TK-deficient HSV-1 infection. We generated TK-deficient HSV-1 by the CRISPR/Cas9 technology. We infected HCFC1 and HCFC1R1 knockout cells with the TK-deficient viruses, and found that both these cells were resistant to TK-deficient viruses (Figure 8E-F). Thus, the deletion of HCFC1 or HCFC1R1 exhibits a strong inhibitory effect on both the wild-type and TK-deficient HSV-1 infection. Our data suggest that HCFC1 and HCFC1R1 may be used as the new drug targets for anti-HSV-1 infection.

## DISCUSSION

To block the virus infection and develop antiviral drugs, both viral components and host factors can be targeted. For viral components, the TK protein of HSV-1 is an effective antiviral target. Acyclovir, a guanosine analogue that targets the TK protein of HSV-1, is the first specific and selective anti-HSV-1 drug(33). However, Acyclovir-resistant HSV-1 strains can lead to severe disease, including disseminated infection of immune-dysregulated individuals (34, 35). Besides, approximately 6-10% of patients co-infected with HSV-1 and HIV are resistant to available anti-herpetic drugs(36). Developing a new, safer and more effective antiviral drug has stimulated intensive efforts in this field.

Alternatively, targeting host factors are another strategy to stop HSV-1 infection. HSV-1 infects cells to complete their genome replication and assembly of viral particles, resulting in lytic death of host cells to release new viruses. Indeed, many proteins in the host cells are essential for HSV-1 infection, reproduction, and lurk. Among them, Nectin-1, HVEM, PILRa and 3-O-S transferases are all generally regarded as HSV-1 receptors expressed on the cell surface(37). Although the membrane receptors are potential targets of HSV-1 infection, their deletions do not completely prevent HSV-1 infection. In addition, due to the capricious mutation, multiple immune escape mechanisms and complex life cycle of HSV-1 virus, it is difficult for the host to eradicate the HSV-1 virus completely (38–40). Targeting multiple pathways is an ultimate choice to prevent HSV-1 infection.

Nectin-1 is recognized as a receptor for HSV-1, and the Nectin-1 knockout cells were to the HSV-1 viruses. However, since the discovery of Nectin-1 in 1998, no effective antiviral drug has been developed by targeting Nectin-1. Here, we compared the antiviral effect of HCFC1-KO, HCFC1R1-KO and Nectin-1-KO cells, and found their resistance efficiency to the HSV-1virus was comparable. Remarkably, the deletion of HCFC1 or HCFC1R1 exhibits a strong inhibitory effect on both wild-type and TK-deficient HSV-1. Therefore, HCFC1 and HCFC1R1 may be used as the new therapeutic targets for fighting infection of both wild-type and TK-deficient HSV-1.

HCFC1 can form a complex with HSV-1 VP16 protein. Our data showed that when the host cell knocks out HCFC1, the nuclear transportation of VP16 protein is hindered, therefore the expression of immediate early genes cannot be activated, and the proliferation of the virus is inhibited. We also showed that loss of HCFC1 potently inhibited viral proliferation. Our data not only confirm that forming the HCFC1 and VP16 complex is important for HSV infection, but also suggest that by directly targeting HCFC1 or by disturbing the HCFC1 and VP16 complex formation can be a useful strategy for inhibiting HSV-1 proliferation.

HCFC1 plays a crucial role in cell cycle regulation and interacts with many proteins, but it remains unclear whether these interactions can play a role in regulating HSV-1 infection (41–43). HCFC1R1 interacts with HCFC1 and is a major regulator of HCFC1. More importantly, HCFC1R1 contains a leucine-rich nuclear export signal and a CRM1-mediated nuclear export signal, suggesting its role in nuclear transportation. Indeed, we demonstrate that the HCFC1R1 deletion blocks both HCFC1 and VP16 nuclear transportation leading to their aggregation in the cytoplasm, thereby inhibits HSV-1 proliferation. Since neither HCFC1 nor HCFC1R1 is a cell membrane receptor and they do not sit on the cell membrane, thus except the cell membrane proteins such as receptors, targeting a host cytosolic factor can also be an effective way to fight HSV-1 infection. In addition, our data showed that the expression of immediate early proteins ICP0 and ICP4, early proteins TK, late proteins VP16 and gD were all affected by the deletion of HCFC1 or HCFC1R1, suggesting that the interaction between HCFC1 or HCFC1R1 and the virus occurs early in the infection stage.

In summary, HCFC1R1, as a regulator of HCFC1, can recognize and form complexes in the early stage of HSV-1 infection that are key to the introduction of HSV-1’s VP16 protein into the nucleus. The HCFC1R1/HCFC1 complex introduces the VP16 protein into the nucleus of the host cell through the nuclear localization signal. VP16 binds with the transcription factor oct-1 and initiates HSV-1 immediate early gene expression. The deletion of HCFC1 or HCFC1R1 causes the blockage of VP16 nuclear transportation, thus interrupts the physiological and metabolic activities of HSV-1. Our data not only provide a mechanism of HSV-1 invading host cells, but also suggest that to develop the inhibitors to interrupt the interaction between HCFC1/ HCFC1R1 complex with VP16 protein can be an effective strategy for treating HSV-1 and its TK mutant strains.

## MATERIALS AND METHODS

### Cell culture

BGC-823 (human gastric carcinoma cells) cells were cultured in RPMI-1640 medium (Hyclone, USA) supplemented with 10% fetal bovine serum (FBS, Hyclone, USA), 100 U/mL penicillin, 100/mL streptomycin (Hyclone, USA) in an incubator at 37°C and 5% CO_2_. Before transfection, the cells were seeded in 6 cm cell culture plates at a cell density of 10^5^. Cell transfection was done using Lipofectamine 3000 reagent (Invitrogen, Carlsbad, CA) according to the manufacturer’s instructions.

### Designing sgRNAs and genetic editing of BGC-823 cells

The sgRNA sequences targeting HCFC1, HCFC1R1, Nectin-1, HSV-1-TK, and HSV-1-VP26 gene loci were selected and designed by using the website http://crispr.mit.edu/. The sgRNA was inserted into the BbsI restriction enzyme site in pX459 plasmid vector. For sgRNA annealing: 1 μl sense and 1 μl anti-sense primers of 100 μM each were mixed with 2 μl 10X Taq polymerase PCR buffer (Takara, Japan) and 16 μl ultra-pure water to a final volume of 20 μl, and subjected to a annealing process to enable hetero-duplex formation at the condition as follows: 95°C, 5 min, 95°C to 85°C at −2.5°C/s, 85°C to 25°C at −0.25°C/s, 25°C, 5 min. Lipofectamine was used to transfect the knock-out plasmids into BGC-823 cells and the cells were screened using puromycin after 24 hours. The cell clones were isolated once the control cells were completely dead. The following primers were used; HCFC1-sgRNA1-5’ -CCCGCCGTAGATCACCAGCT-3’ for BGC-823 cells; HCFC1R1-sgRNA1-5’-TTTCCCAGCTCCCCTCTCCG-3’ for BGC-823 cells; 5’ -TCTGGAGGTAGTCCCAAGCC-3’ and 5’ -GAGGAAAGCACGAAGTGGTATG-3’ for YZ-BGC-823-HCFC1; 5’ -GCGTGAGCTGAGATGGGACT-3’ and 5’ -GTGAGGACCGCCTGTGATTC-3’ for YZ-BGC-823-HCFC1R1.

### Protein extraction, Western blot analyses and antibodies

The cells were washed with ice-cold PBS, harvested by gentle scraping, and lysed with the protein extraction buffer containing 150 mM NaCl, 10 mM Tris (pH 7.2), 5 mM EDTA, 0.1% Triton X-100, 5% glycerol, and 2% SDS. Protein concentrations were determined by BCA Protein Assay Kit (P0010S, Beyotime, USA). Forty μg of total proteins were separated by electrophoresis on 10% polyacrylamide gels and transferred to PVDF membranes (Bio-Rad, CA, USA). The membranes were blocked with 5% BSA in Tris-buffered saline for 1 h at room temperature. After overnight incubation at 4°C with primary antibodies, the antigens were detected by IRDye 800CW secondary antibody (1:10000) and visualized by Odyssey Infrared Imaging System (LI-COR) (Westburg, Netherlands). Primary antibodies used for Western blotting were HCFC1 rabbit antibody (Cell Signaling, USA), HCFC1R1 rabbit mAb (Cell Signaling, USA), VP16 mouse monoclonal antibody (Proteintect, China) and anti-GAPDH (Abcam, England); Secondary antibodies were rabbit anti-mouse IgG H&L (HRP), goat anti-rabbit IgG H&L (HRP) (Abcam, England). The expression of β-actin was used as control.

### Attachment assay

The cells were seeded in 6 cm cell culture plates at a cell density of 2 × 10^5^. Before infection, HSV-1 was pretreated with 2 μg/ml DNase (Takara, Japan), and then diluted to MOI=20, 50, 100 in DMEM medium (Hyclon, USA). The pre-cooling HSV-1 was added to wild-type and BGC-823^HCFC1R1−/−^ cells for 1 h at 4°C, respectively. The cells were washed three times with PBS, and then the genomic DNA was isolated by EasyPure Genomic DNA Kit (Transgen, China). The copy number of HSV-1 binding was examined by real-time quantitative polymerase chain reaction (qPCR).

### Quantitative PCR

The HSV-1 copy number was examined by qPCR using SYBR/ROX (RR82LR; Takara, Japan) on ABI qPCR machine (Lifetech, CA, USA). The ΔΔCT values of qPCR were calculated using the manufacturer’s software and used to estimate the levels of HSV-1 genomic DNA. qPCR primers were following: qPCR-HSV-1-gD-F 5’-CGCCGTCAGCGAGGATAA-3’, qPCR-HSV-1-gD-R 5’TCTTCACGAGCCGCAGGTA-3’; qPCR-HSV-1-gB-F 5’GTCGG CAAGGTGGTGATGG-3’, qPCR-HSV-1-gB-R 5’ GTAGCGAAAGGCGAAGAAGG-3’; qPCR-HSV-1-TK-F 5’ CGATGACTTACTGGCGGGTG-3’, qPCR-HSV-1-TK-R 5’GGTCG AGGCGGTGTTGTGT-3’; qPCR-HSV-1-ICP0-F 5’GTGCATGAAAACCTGGATGC-3’, qPCR-HSV-1-ICP0-R 5’TTGCCCGTCCAGATAAAGTC-3’. Changes on the levels of different samples were evaluated after normalized to the β-actin control.

### Cellular proliferation assays

TransDetect Cell Counting Kit (CCK) (TransGene, Biotech, China)-based measurements of cellular proliferation were performed by plating 2×10^3^ cells per well in 96-well plates. Three replicate wells were plated for each sample with 100 μl medium. At the initial time point or after 24, 48 or 72 hours, 10 μl CCK was added into the cell culture well and then continue to culture for 2 hours. The absorbance at 450 nm was determined by a microplate reader and the cell proliferation rate was counted.

### Immunofluorescence analyses

BGC-823 cells were plated onto coverslips and transfected with various overexpression plasmids as described above. Twenty four hours after transfection, the cells were infected with HSV-1 (moi=100) for indicated time points at 4°C and fixed with 2% paraformaldehyde in PBS for 20 min. After a brief treatment with methanol (5 min), the coverslips were incubated with the primary antibodies against Flag (CST; 1:200), HCFC1 rabbit antibody (Cell Signaling, USA), HCFC1R1 rabbit mAb (Cell Signaling, USA) or VP16 mouse monoclonal antibody (Proteintect, China) in PBS plus 2% BSA overnight at 4°C in a humid chamber. The next day, the coverslips were incubated with Alexa Fluor 488–conjugated goat antibody to rat IgG (A-11006; Molecular Probes; 1:200), Alexa Fluor 594–conjugated goat antibody to rat IgG (A-11006; Molecular Probes; 1:200), Alexa Fluor 488–conjugated goat antibody to mouse IgG (A-11001; Molecular Probes;1:200) or Alexa Fluor 594–conjugated goat antibody to mouse IgG (A-11006; Molecular Probes; 1:200) for 2 h at room temperature. Images were collected by using a Zeiss LSM700 confocal microscope.

### Statistical Analysis

For comparisons between two groups, statistical analyses were done by the Student’s *t*-test (unpaired and two-tailed) using Prism 7.0c (GraphPad). All experiments were performed repeatedly at least three times. Error bars represent standard deviations (SDs). Findings were considered to be significantly different when *p* < 0.05.

## ACKNOWLEDGMENTS

This study was supported by Natural Science Foundation of the Fujian Province, China (Grant No.2017J01621), Innovative Research Teams Program II of Fujian Normal University in China (IRTL1703), Fujian Key Laboratories Funds, and a Fujian Provincial Lingjun Scholarship to QC. We would like to thank Dr. Lijun Sun for his valuable advises and suggestions, Shaoli Cai and Zhang Lin for administrative assistance, and Dr. Daliang Li and the members of the Chen’s laboratory for technical assistance and helpful discussion.

